# Transmission potential of *Culex* and *Aedes* species for Madariaga virus, a member of the eastern equine encephalitis virus complex

**DOI:** 10.1101/2025.09.02.673670

**Authors:** Danilo de Carvalho-Leandro, Francisco C. Ferreira, Nadia Fernandez Santos, William J. Sames, Yuexun Tian, Martial L. Ndeffo-Mbah, Erica A. Costa, Michael J. Turell, Gabriel L. Hamer, Tereza Magalhaes

**Affiliations:** Department of Entomology, Texas A&M University. College Station-TX, 77843, United States of America (USA); Colégio de Aplicação, Universidade Federal de Pernambuco. Recife-PE, 50670, Brazil; Instituto Politecnico Nacional, Centro de Biotecnologia Genomica. Reynosa, Tamaulipas 88710, Mexico; PO Box 547, Leakey-TX, 78873, USA; Department of Integrative Biosciences, College of Veterinary Medicine and Biomedical Sciences, Texas A&M University, College Station-TX, 77843, USA; Research Laboratory in Animal Virology, Veterinary School, Universidade Federal de Minas Gerais. Belo Horizonte-MG, 31270, Brazil; VectorID LLC, Frederick-MD, 21702, USA

**Keywords:** Vector competence, South American eastern equine encephalitis virus, Madariaga virus, alphavirus, equine encephalomyelitis, equine encephalitis, *Aedes taeniorhynchus*, *Aedes albopictus*, *Aedes aegypti*, *Culex tarsalis*, *Culex coronator*, *Culex quinquefasciatus*, vector-borne disease, mosquito-borne disease, mosquito

## Abstract

Madariaga virus (MADV), widely distributed in Latin America, can cause severe disease in humans and equids, yet, key aspects of its transmission cycle remain unclear. To identify mosquitoes that could act as MADV vectors, we assessed the vector competence of *Aedes aegypti*, *Ae. albopictus*, *Ae. taeniorhynchus*, *Culex tarsalis*, *Cx. coronator*, and *Cx. quinquefasciatus*, following oral exposure to MADV isolated in Panama (all species) or Brazil (*Ae. taeniorhynchus* only). MADV RNA and infectious virus were quantified from mosquito bodies, legs, and saliva. At 14 days post-exposure, five species had virus in all biological sample types. *Culex quinquefasciatus* was susceptible to infection and dissemination but had no positive saliva samples. *Aedes taeniorhynchus* showed higher infection probabilities with MADV-BR. Time-course analysis revealed distinct dynamics in *Ae. aegypti* and *Ae. albopictus*. Our findings indicate MADV may be compatible with mosquito species present in endemic regions and areas at risk of virus introduction.

## 1. Introduction

Madariaga virus (MADV), an encephalitic alphavirus belonging to the eastern equine encephalitis virus (EEEV) complex, is prevalent in Latin America and a potential emerging pathogen (1). The virus circulates in enzootic cycles involving sylvatic mosquito vectors and vertebrate animals and can cause disease in equids and humans during spillover events. Case fatality rates among equines can reach >90% during epizootics (1), and severe and fatal human cases due to MADV infection have been documented (2–6).

The vectors of MADV remain unknown, and different mosquito species likely act as vectors across the various biomes where the virus circulates in Latin America (1). Nonetheless, field and laboratory data have helped identify potential candidates. In the field, MADV has been isolated from several mosquito species, with isolations from *Culex* (*Melanoconion*) spp. being more frequent (1, 7–12). The virus has also been detected in *Aedes taeniorhynchus* on more than one occasion, including during an equine epizootic in Brazil in 1960 (11, 13), suggesting this species may act as an amplification and bridge vector during outbreaks in certain regions. A limitation in these field studies is the lack of information about the physiological status of mosquitoes (i.e., whether females were engorged or not), raising the possibility that virus detection could have been from an infected host’s blood and not from mosquito infection.

As for laboratory studies, to date only one vector competence study has been published on MADV. In that study, mosquito species collected in the Peruvian Amazon were tested for their ability to transmit the virus using a live animal model (14). The following species were shown to be competent vectors: *Cx.* (*Mel.*) *pedroi*, *Ae.* (*Ochlerotatus*) *fulvus*, *Psorophora* (*Janthinosoma*) *albigenu*, *Ps.* (*Grabhamia*) *cingulata*, *Ae.* (*Och.*) *serratus*, and *Ps.* (*Jan.*) *ferox*.

Given the limited data on the identification of MADV vectors, we assessed the ability of selected mosquito species – most of which had not been previously tested – to transmit the virus following oral exposure. Species selection included one or more of the following criteria: 1) high abundance across various geographic regions within the Americas; 2) known role as a vector of arboviruses; 3) feeding habit on mammalian hosts indicating ability to act as bridge vectors; 4) evidence for having a potential role in enzootic or epizootic MADV transmission; and 5) experimental feasibility. The species tested were *Cx. coronator*, *Cx. quinquefasciatus*, *Cx. tarsalis*, *Ae. aegypti*, *Ae. albopictus*, and *Ae. taeniorhynchus*.

## 2. Methods

### 2.1 Mosquitoes

The vector competence for MADV was evaluated in six mosquito species: *Aedes aegypti*, *Ae. albopictus*, *Ae. taeniorhynchus*, *Culex coronator*, *Cx. tarsalis*, and *Cx. quinquefasciatus*. *Aedes aegypti* and *Ae. albopictus* eggs were collected in September 2023 in Hidalgo County and Brazos County, Texas, respectively, using ovitraps. Adult *Ae. taeniorhynchus* were collected in August 2024 in Hidalgo County, Texas, using modified CDC light traps. *Culex coronator* larvae were collected in October 2024 in Leakey, Texas, using a larval dipper. *Culex tarsalis* and *Cx. quinquefasciatus* and were obtained from established colonies maintained at Colorado State University. All specimens were transported to insectaries at Texas A&M University and reared under controlled conditions (28°C, 70% humidity) with *ad libitum* access to 10% sucrose and water. Adults of *Ae. taeniorhynchus* and *Cx. coronator* parental generation (F0), and F2-F3 adults of *Ae. aegypti* and *Ae. albopictus* were used in the experiments. *Culex tarsalis* and *Cx. quinquefasciatus* adult females used in the experiments were of unknown generation. All mosquitoes were 4-6 days old at the time of virus exposure, except for *Ae. taeniorhynchus*, which were collected as adults from the field and used directly in the experiments, therefore these were of unknown age.

Mosquito specimens brought from the field (F0) and the first filial generation (F1) were reared in the Arthropod Containment Level (ACL)-2 room; after that, mosquitoes were reared in ACL-1 rooms.

### 2.2 Viruses

MADV-PAN (Lineage III; GenBank: KJ469648), isolated from a horse during a 2010 outbreak in Panama, was provided by the World Reference Center for Viruses and Arboviruses at the University of Texas Medical Branch, and used to expose all mosquito species. Passage 3 MADV-PAN was received lyophilized and subsequently resuspended in cell culture medium. MADV-BR (Lineage III; GenBank: MZ389692), isolated by our group in 2019 from the central nervous system of a horse during an epizootic in Northeast Brazil (15), was used only to expose *Ae. taeniorhynchus*. Passage 9 MADV-BR was received as frozen aliquots. The two strains share 95% nucleotide and 99% protein identity.

MADV-PAN and MADV-BR were propagated in Vero cells (CCL-81, ATCC) maintained at 37°C with 5% CO_2_ in 5% complete Dulbecco’s Modified Eagle Medium (DMEM). Virus stocks were generated by infecting ∼80% confluent monolayers and harvesting cells and supernatant at 36 h post-infection, when cytopathic effect reached 80-90%. The collected material was centrifuged (1,000xg, 5 min, 4°C), and the supernatant was aliquoted and stored at -80°C. Final stock titers were 1.6 x 10^7^ PFU/mL for MADV-PAN (passage 5) and 4.0 x 10^7^ PFU/mL for MADV-BR (passage 11). For mosquito infections, viruses were propagated using the same methodology, but the supernatant was mixed with blood immediately after centrifugation.

All procedures involving MADV were conducted in Biosafety Level 3 and ACL-3 laboratories under approved protocol IBC2022-075.

### 2.3 Mosquito exposure

Adult female mosquitoes were transferred to the ACL-3 and acclimated for 1-2 days in a growth chamber (28°C, 70% humidity). Groups of 200-400 mosquitoes were starved for 24 h, then offered a 1:1 mixture of defibrinated calf blood and MADV-PAN or MADV-BR via membrane feeder. After ∼30 minutes, engorged females were sorted, transferred to 473 mm^3^-cartons, and maintained with 10% sucrose and water for up to 21 days. An aliquot of the blood:virus mixture was kept at 37°C during feeding and later stored at -80°C for bloodmeal titration. Two experimental replicates were performed per mosquito species, and per *Ae. taeniorhynchus* exposure with each virus strain.

### 2.4 Sample collection

Thirteen to 30 MADV-exposed mosquitoes per group – and per time point for *Ae. aegypti* and *Ae. albopictus* – were sampled in each experimental replicate. Bodies, legs, and saliva were collected for RNA and infectious virus quantification. Collections were performed at 14 days post-exposure (dpe) for all species, and additionally at 3, 7, and 21 days for *Ae. aegypti* and *Ae. albopictus*. For sample collection, females were anesthetized (∼2°C), and legs were removed and placed in microcentrifuge tubes with mosquito diluent (PBS with 20% FBS, antibiotics, antimycotic) and borosilicate beads.

Wings were discarded. Saliva was collected at room temperature by forced salivation, by inserting the proboscis into glass capillaries containing immersion oil. After 30 min of salivation, capillaries were transferred to microcentrifuge tubes with mosquito diluent and centrifuged (11,000xg, 3 min, 2°C).

Mosquito bodies were then placed in microcentrifuge tubes with mosquito diluent and borosilicate beads. Leg and body samples were homogenized with a TissueLyzer II. All samples were stored at - 80°C. For molecular and plaque assays, samples were thawed, centrifuged (11,000xg, 3 min, 2°C), and supernatants used.

### 2.5 RNA extraction and reverse transcription quantitative PCR (RT-qPCR)

RNA was extracted from virus stocks, bloodmeal samples, and mosquito biological samples using the Mag-Bind™ Viral DNA/RNA Kit (Omega Bio-tek) on a KingFisher™ Flex System with 96-deep well head, following the manufacturers’ instructions. A 50-µL sample volume was used, and RNA was eluted in ultra-pure water. For RNA extraction, virus stocks were first serially diluted in 10-fold dilutions (undiluted to 10^-9^) in 5% DMEM, and bloodmeal and mosquito samples were processed after stored material was thawed and centrifuged.

The following primers and probe targeting MADV nsP2 (nt 1776-1892) were designed using IDT PrimerQuest™: forward (5’-3’) – GGCTGAACAGGTGCTAGTTAT; reverse (5’-3’) – CTATTCCAATCCCGGACTTTCA; probe (5’-3’) – CGCGCCGGTAGGTACAAAGTAGAA (6-

FAM/ZEN™/3’ IB™ FQ). Amplicon size was 117 bp. Reactions were prepared using the iTaq Universal Probes One-Step Kit (Bio-Rad). Thermocycling was performed on a Bio-Rad CFX96 under the following conditions: 50°C for 10 min, 95°C for 2 min; 40 cycles of 95°C for 15 sec and 60°C for 30 sec. Each run included a MADV RNA standard curve (10^-2^-10^-6^) and a no-template (water) control. A cycle threshold (Ct) ≤38 was considered positive.

### 2.6 Plaque assays

To confirm the presence of infectious virus, all saliva and ∼50% of body and leg RT-qPCR-positive samples were tested by plaque assay. Virus stocks and bloodmeal contents were also assayed. For that, Vero cells in 24-well plates were inoculated with samples, followed by 1 h adsorption and addition of overlay (1:1 Tragacanth medium: 2x 8% complete DMEM). Plaques were visualized after monolayer staining with 0.1% crystal violet in 20% ethanol and counted to calculate PFU/mL. Bloodmeal samples were plated in duplicate for each dilution (10^-2^-10^-6^); body and leg samples were plated in one replicate per dilution (10^-2^ and 10^-3^); and saliva samples (post-RNA extraction, ∼20 µL) were plated at a 10^-2^ dilution in a single well. Negative controls (medium only) were included in each plate. Fifteen RT-qPCR-negative samples per tissue type were randomly selected and also assayed.

### 2.7 Data analysis

Infection, dissemination, and transmission rates across mosquito species were calculated as the proportion of RT-qPCR-positive samples per body, leg, and saliva samples, respectively. No statistical tests were applied here, as differences were assessed using regression models.

For more robust statistical analysis and to account for blood meal titers and replicates, infection status probabilities per mosquito biological sample (note that this is different than ‘infection rate’, which is based on midgut data only) were estimated using logistic regression models with fixed effects and covariates (‘blood meal titer’ and ‘replicate’, retained if significant). For that, GLIMMIX procedure was used in SAS Studio^®^. Least-square means, 95% confidence intervals (CIs), and odds ratios (ORs) were obtained. Pairwise comparisons were conducted with ORs with significance set at p<0.05.

Models were applied to: (i) compare MADV-PAN infection probabilities across species (per biological sample type) at 14 dpe (‘mosquito species’ as fixed effect; ref: *Ae. aegypti*); (ii) compare MADV-PAN and MADV-BR infection probabilities per biological sample type in *Ae. taeniorhynchus* at 14 dpe (‘virus strain’ as fixed effect; ref: MADV-PAN); and (iii) assess the temporal infection dynamics of MADV-PAN in *Ae. aegypti* and *Ae. albopictus* (‘mosquito species’, ‘dpe’, and ‘mosquito species x dpe’ interaction; refs: *Ae. aegypti*, 3 dpe).

In (i), groups without positive saliva samples were excluded and compared using Fisher’s exact tests with FDR and Bonferroni correction. In (iii), time points with zero positives were retained to preserve model structure.

Virus titers (PFU/mL) were log_10_-transformed and plotted for visualization. Graphs were generated in GraphPad Prism 10.5.0.

## 3. Results

### 3.1 MADV-PAN infection, dissemination, and transmission rates across mosquito species

Bloodmeal titers for MADV-PAN ranged from 1.1 x 10^5^ to 5.8 x 10^7^ PFU/mL. All six species were susceptible to infection (body) and dissemination (legs). Virus was detected in the saliva of all species, except *Cx. quinquefasciatus* (Table 1). *Aedes aegypti* had the highest infection rate (53%, 84/160).

**Table 1:**
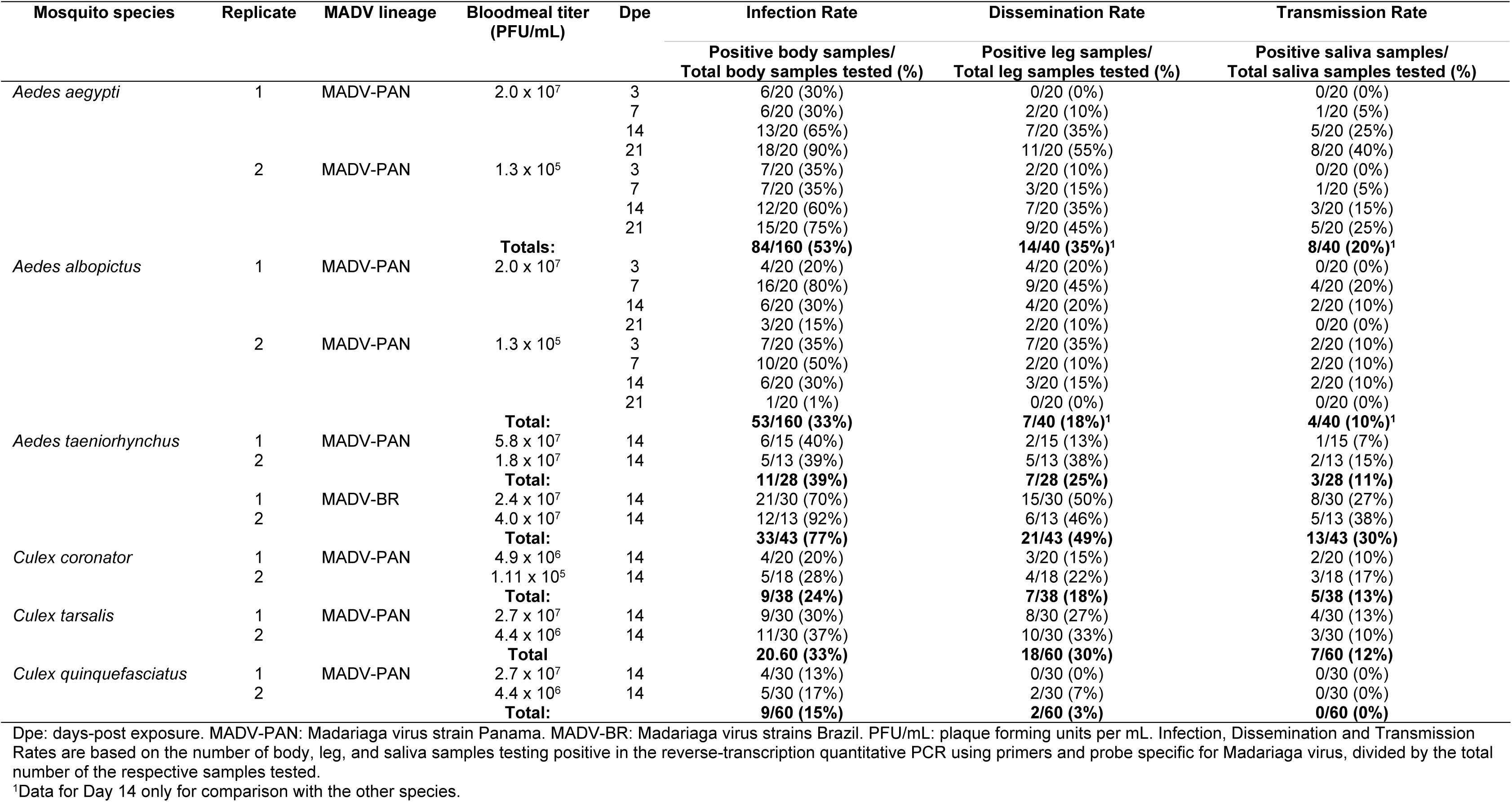
Numbers of mosquito body, legs, and saliva samples that tested positive through reverse transcription quantitative PCR using Madariaga virus-specific primers and probe.

Temporal trends were examined in *Ae. aegypti* and *Ae. albopictus*. In *Ae. aegypti*, dissemination and transmission rates increased over time, reaching 50% and 33% at 21 days post-exposure (dpe), respectively. In contrast, *Ae. albopictus* showed decreasing rates after 7 dpe, with dissemination falling to 5% and transmission to 0% by 21 dpe (Table 1).

### 3.2 MADV-PAN infection probabilities in body, legs, and saliva across mosquito species

Predicted infection probabilities in bodies and legs were adjusted for bloodmeal titer; replicate was excluded. Mosquito species significantly predicted infection status in both body (p<0.001) and leg (p=0.008) models.

*Aedes aegypti* had the highest predicted body infection probability, followed by *Ae. taeniorhynchus*, *Cx. tarsalis*, *Ae. albopictus*, *Cx. coronator*, and *Cx. quinquefasciatus* (Fig. 1A, Supp. Table 1). Significant pairwise differences were found between *Ae. aegypti* and all other species, and between *Cx. quinquefasciatus* and several species (Supp. Table 2).

**Fig. 1:**
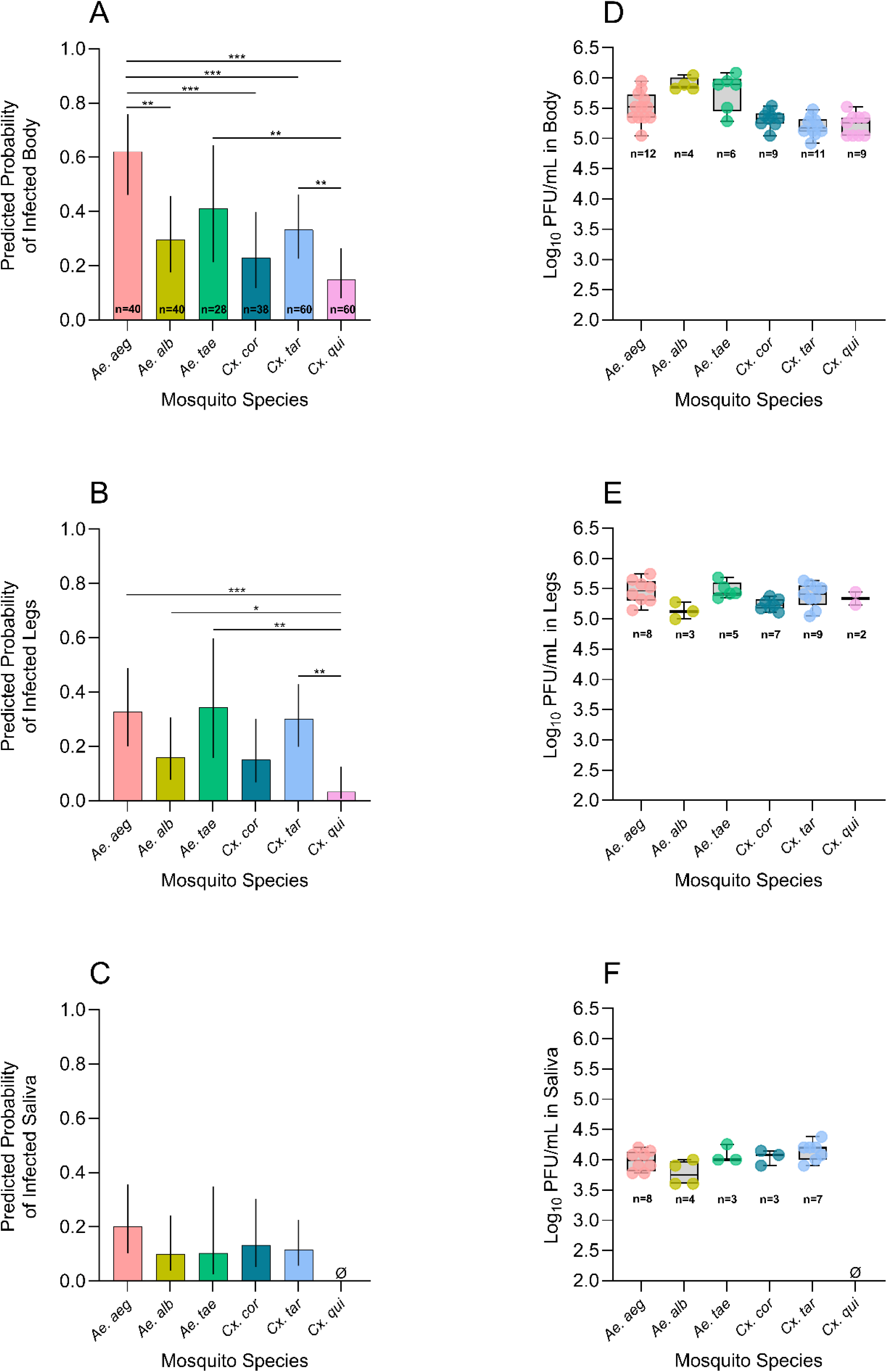
Predicted infection probabilities of Madariaga virus strain Panama (MADV-PAN) derived from regression models in body (A), legs (B), and saliva (C), and log_10_-transformed MADV-PAN titers in body (D), legs (E), and saliva (F) samples collected from distinct mosquito species at 14 days-post exposure. In A-C, models were adjusted for blood meal titers, and two experimental replicates per mosquito species were accounted for; the total number of mosquitoes dissected per species across both replicates is shown in panel A, and vertical lines indicate 95% confidence intervals. In D-E, the number of samples tested is shown below each group, and box and whisker plots indicate the median and interquartile range – all of the saliva samples and approximately 50% of body and legs testing positive in the molecular assay were tested in plaque assays. *Ae. aeg*: *Aedes aegypti*; *Ae. alb*: *Aedes albopictus*; *Ae. tae*: *Aedes taeniorhynchus*; *Cx. cor*: *Culex coronator*; *Cx. tar*: *Culex tarsalis*; *Cx. qui*: *Culex quinquefasciatus*. In (C), data from Cx. *quinquefasciatus* was removed from the model as this species had zero positive saliva samples, leading to complete data separation. In (F), *Cx. quinquefasciatus* data is absent as the model was fit only with qRTPCR-positive data. Bars with asterisks in A-C indicate the level of statistical significance between the two groups shown at the end of each bar, based on odds ratio pairwise comparisons: *p<0.05, **p<0.001, ***p<0.0001. PFU/mL: plaque forming units per mL.

Leg infection probabilities were highest in *Ae. aegypti*, *Ae. taeniorhynchus*, and *Cx. tarsalis*, and lowest in *Cx. quinquefasciatus* (Fig. 1B, Supp. Table 1). Odds ratios showed significant differences between *Cx. quinquefasciatus* and four other species (Supp. Table 2).

In the saliva model, *Cx. quinquefasciatus* was excluded due to lack of positives. Species was not a significant predictor of the outcome, though *Ae. aegypti* showed a non-significant trend toward higher infection probabilities (Fig. 1C, Supp. Table 1). Fisher’s exact tests showed that *Cx. quinquefasciatus* differed significantly from two or all other species based on Bonferroni or FDR corrections, respectively (Supp. Table 3).

### 3.3 MADV-PAN and MADV-BR infection probabilities in Ae. taeniorhynchus

Overall, *Ae. taeniorhynchus* exposed to MADV-BR showed a trend for higher predicted infection probabilities in body, leg, and saliva samples compared to those exposed to MADV-PAN, although statistically significant difference was only observed in body samples (Fig. 2A-C, Supp. Tables 4 and 5). Models were adjusted for bloodmeal titer; replicate was excluded.

**Fig. 2:**
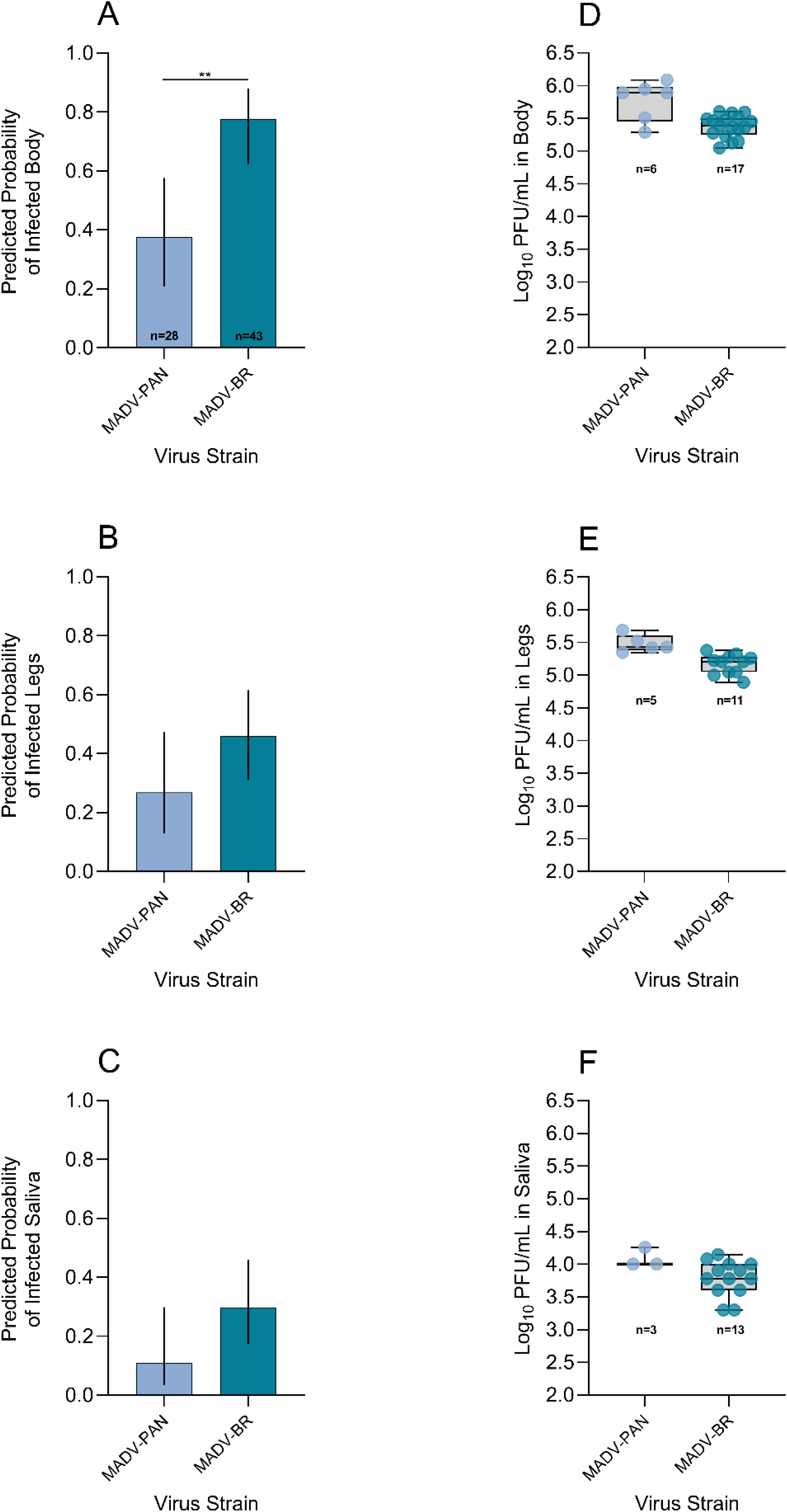
Predicted infection probabilities of Madariaga virus strain Panama (MADV-PAN) and Madariaga virus strain Brazil (MADV-BR) derived from regression models in body (A), legs (B), and saliva (C), and log_10_-transformed MADV-PAN and MADV-BR titers in body (D), legs (E), and saliva (F) samples collected from *Aedes taeniorhynchus* at 14 days-post exposure. In A-C, models were adjusted for blood meal titers, and two experimental replicates per virus strain were accounted for; the total number of mosquitoes dissected per species across both replicates is shown in panel A, and vertical lines indicate 95% confidence intervals. In D-E, the number of samples tested is shown below each group, and box and whisker plots indicate the median and interquartile range – all of the saliva samples and approximately 50% of body and legs testing positive in the molecular assay were tested in the plaque assay. Bars with asterisks in A-C indicate the level of statistical significance between the two groups shown at the end of each bar, based on odds ratio pairwise comparisons: *p<0.05, **p<0.001, ***p<0.0001. PFU/mL: plaque forming units per mL.

### 3.4 Temporal MADV-PAN infection probability in Ae. aegypti and Ae. albopictus body, legs and saliva

In the body model (adjusted for bloodmeal titer), mosquito species (p<0.001) and mosquito species x dpe interaction (p<0.0001) were significant predictors of the outcome, while dpe alone was not (p=0.1). In *Ae. aegypti*, predicted body infection probabilities increased over time. In *Ae. albopictus*, they rose from 3 to 7 dpe and declined thereafter. Significant differences between species were observed at 7, 14, and 21 dpe (Fig. 3A, Supp. Tables 6 and 7).

**Fig. 3:**
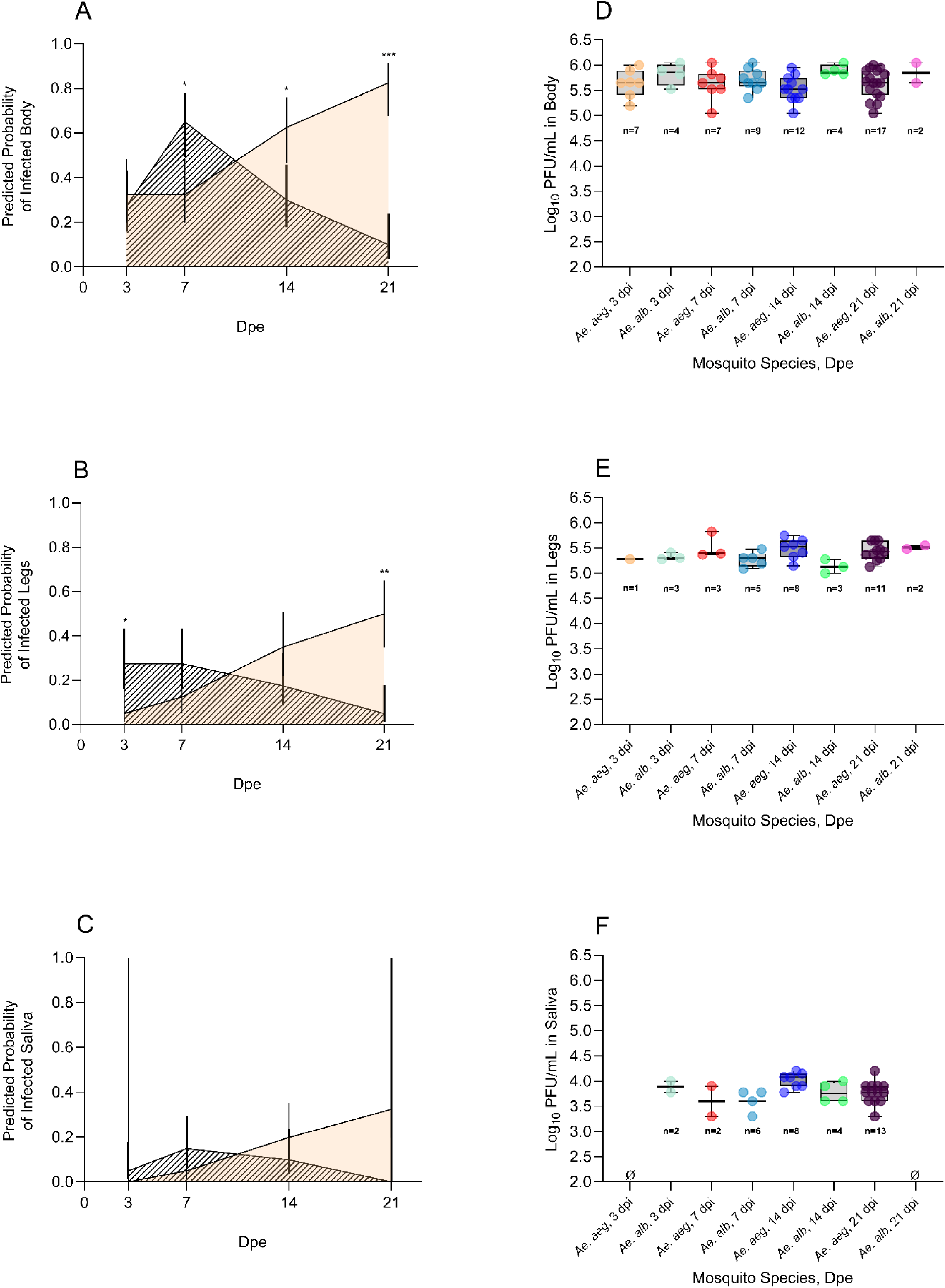
Predicted infection probabilities of Madariaga virus strain Panama (MADV-PAN) derived from regression models in body (A), legs (B), and saliva (C), and log10-transformed MADV-PAN titers in body (D), legs (E), and saliva (F) samples collected from Aedes aegypti (Ae. aeg; orange shading) and Aedes albopictus (Ae. alb; diagonal line shading) at 3, 7, 14, and 21 days-post exposure (dpe). In A-C, 20 individual mosquitoes were dissected per species per time point, and models were adjusted for blood meal titers, and two experimental replicates per virus strain were accounted for; vertical lines indicate 95% confidence intervals, asterisks indicate the level of statistical significance between the two groups at a given time point, based on odds ratio pairwise comparisons: *p<0.05, **p<0.001, ***p<0.0001. The wide confidence intervals in (C) reflect the absence of positive saliva samples in Ae. aegypti at 3 dpe and Ae. albopictus at 21 dpe – these groups were retained because the model included a ‘mosquito x dpe’ interaction, and variation at other time points allowed estimation. In D-F, box and whisker plots indicate the median and interquartile range; the number of samples tested is shown below each group – all of the saliva samples and approximately 50% of body and legs testing positive in the molecular assay were tested in the plaque assay. PFU/mL: plaque forming units per mL.

In the leg model, only mosquito species x dpe interaction was significant (p< 0.0001). Trends were similar to those in body samples, with increasing infection probabilities over time in *Ae. aegypti*, and a peak at 7 dpe followed by a decline in subsequent time points in *Ae. albopictus*. Significant differences between the two species were observed at 3 and 21 dpe (Fig. 3B, Supp. Tables 6 and 7).

In the saliva model, no fixed effect was a predictor of the outcome, but trends between *Ae. aegypti* and *Ae. albopictus* were similar to those of body and legs (Fig. 3C, Supp. Tables 6 and 7).

### 3.5 MADV-PAN titers in body, legs, and saliva across mosquito species

All RT-qPCR-positive saliva, body and leg samples tested by plaque assay yielded plaques, except for one *Ae. albopictus* saliva sample (7 dpe, replicate 2). None of the 45 RT-qPCR-negative samples tested yielded plaques.

Mean MADV-PAN titers ranged from 5.2-5.9 and 5.1-5.5 log_10_ PFU/mL in bodies and legs, respectively (Fig. 1D-E, Supp. Table 8). Saliva PFU counts ranged from 1 to 12 per well at a 10^-2^ dilution. Given the volume used for RNA extraction and additional volume loss, these values derive from <1 µL of actual saliva, with resulting titers of 3.7-4.3 log_10_ PFU/mL across competent species (Fig. 1F, Supp. Table 8).

### 3.6 MADV-PAN and MADV-BR titers in Ae. taeniorhynchus body, legs and saliva

Mean log_10_-transformed MADV-PAN and MADV-BR titers in *Ae. taeniorhynchus* body samples were 5.8 and 5.3, respectively (Fig. 2D, Supp. Table 9). In legs, titers were 5.5 and 5.2 for MADV-PAN and MADV-BR, respectively (Fig. 2E, Supp. Table 9). In saliva, titers were 4.1 and 3.8 for MADV-PAN and MADV-BR, respectively (Fig. 2F, Supp. Table 9).

### 3.7 MADV-PAN titers in Ae. aegypti and Ae. albopictus over time

In RT-qPCR-positive samples, mean body titers ranged from 5.5 (14 dpe, *Ae. aegypti*) to 6.0 (21 dpe, *Ae. albopictus*) log_10_ PFU/mL (Fig. 3D, Supp. Table 10). Leg titers ranged from 5.1 (*Ae. albopictus*, 14 dpe) to 5.6 (*Ae. aegypti*, 7 dpe) log_10_ PFU/mL (Fig. 3E, Supp. Table 10). Saliva titers ranged from 2.8 (*Ae. albopictus*, 7 dpe) to 4.2 (*Ae. albopictus*, 3 dpe) log_10_ PFU/mL (Fig. 3F, Supp. Table 10).

## 4. Discussion

Five of the six mosquito species tested had MADV RNA and infectious particles in their saliva following oral exposure indicating that MADV may be compatible with and potentially transmitted by a broad range of mosquito vectors in the field. As expected for arbovirus infections, infection rates declined from body to legs to saliva, reflecting known biological barriers encountered by arboviruses within mosquito tissues (16–18). The agreement between RT-qPCR and plaque assay results (where all but one RT-qPCR-positive sample yielded plaques, while none of the negative samples did) validated our molecular assay.

MADV-PAN body and leg infection probabilities varied across the six mosquito species at 14 dpe, likely reflecting species-specific midgut infection and escape barriers. Conversely, saliva infection probabilities were more uniform among competent species, although, as noted, it is possible that low sample sizes may have masked differences.

The absence of positive saliva samples in *Cx. quinquefasciatus*, seen in both replicates, indicates that this species may not be competent for MADV-PAN. This species also had the lowest infection probabilities in body and legs, suggesting limited ability of the virus to infect and disseminate within its tissues.

Among the competent species, *Cx. tarsalis* is an ornithophilic mosquito that sporadically feeds on mammals. *Culex tarsalis* is a major WNV vector and an important vector for western equine encephalitis in North America (19). Although this species is not found in MADV-endemic areas, its potential to transmit MADV becomes relevant if the virus is introduced in regions within its range, particularly if birds are shown to participate in MADV transmission cycles.

*Culex coronator*, also competent for MADV, is widespread throughout the Americas (20–24). This species has been implicated as a vector of certain arboviruses (25), but it is usually not included in disease surveillance testing. While *Cx. coronator* has generally been associated with sylvatic habitats, there are reports of its presence in urban and peri-urban areas (26, 27). Interestingly, *Cx. coronator* from Peru did not transmit MADV in the study by Turell *et al*. (14) – this discrepancy may be due to differences in mosquito and virus strains used in each study.

*Aedes aegypti*, another MADV-competent species in our study, is a major vector of arboviruses in urban and peri-urban environments due to its strong anthropophilic behavior (28, 29). This behavior makes its involvement in enzootic transmission cycles unlikely. However, this mosquito is also present in rural areas, and host-feeding behavior can vary geographically, with some populations occasionally feeding on non-human mammals (29–31). Given this variability and the species broad distribution throughout the Americas, its potential role as a bridge vector in MADV-endemic regions cannot be entirely ruled out.

MADV infection probabilities in body, legs, and saliva was assessed over time in *Ae. aegypti* and *Ae. albopictus*. *Ae. albopictus* was included in this study due to its global distribution (32, 33), wide host range (29, 32), and potential role as a bridge vector for arboviruses (34). The temporal infection dynamics differed sharply between these species. In *Ae. aegypti*, probabilities of infection, dissemination, and transmission increased over time. In *Ae. albopictus*, infection probabilities increased from 3 to 7 dpe but declined thereafter. Regardless of the mechanisms, which warrants further investigations, the observed infection dynamics of MADV-PAN in these two species highlight the need to account for temporal variation in mosquito-virus interactions when modeling transmission, as they may help explain spatiotemporal heterogeneity in arboviral disease patterns. Overall, infection patterns of arboviruses in mosquito tissues vary in the literature and appear to depend on mosquito and virus species, strains, bloodmeal titers, and environmental factors (35–39).

*Aedes taeniorhynchus*, also called the black salt marsh mosquito, was competent for both MADV-PAN and MADV-BR. While infection probabilities for MADV-PAN were similar to those of other species, probabilities showed a higher trend for MADV-BR. Although not statistically significant in leg and saliva samples, this trend was consistent across biological sample types and replicates, suggesting a true biological difference. This species is widely distributed throughout coastal and some inland regions of the Americas and is capable of exploiting diverse habitats (40–46). It has a broad vertebrate host range (47, 48) and although typically considered a nuisance species, it is competent for several medically important arboviruses (40, 41, 49–58). Although MADV was isolated from *Ae. taeniorhynchus* during an equine epizootic in Brazil in 1960 (59), the transmission potential of this species had not been experimentally confirmed. Our results, combined with the species ecological features and the prior isolation of the virus from field specimens, support its role as a potential amplification and bridge vector during outbreaks.

Overall, the findings in *Ae. aegypti*, *Ae. albopictus* and *Ae. taeniorhynchus* reinforce the need to investigate species- and strain-specific mosquito-virus interactions more thoroughly.

This study has limitations. For instance, while four species were tested using F0-F3 generations to better reflect field conditions, two species (*Cx. tarsalis* and *Cx. quinquefasciatus*) came from long-term established colonies, which may limit the generalizability of the results to wild populations. In addition, the viral titers and artificial oral infection method used may have either over- or under-estimated infection probabilities. Finally, we were unable to include *Culex* (*Melanoconion*) spp. – considered the primary candidates for enzootic MADV transmission – due to colonization difficulties and limited field collections (1, 12). We encourage the inclusion of this group in future vector competence studies with MADV.

Despite these limitations, our study provides new insights into the transmission potential of mosquito species found in the United States, including those broadly distributed in the Americas (e.g., *Ae. taeniorhynchus*) and globally (e.g., *Ae. aegypti* and *Ae. albopictus*), for MADV. Although this pathogen remains poorly studied, it has been associated with high morbidity and mortality rates in equines and with severe disease and death in humans (1). Identifying mosquito species able of transmitting MADV is essential for anticipating potential emergence events, particularly as the virus expands geographically. Studies like ours are critical for understanding arbovirus emergence and for risk assessment, more so when integrated with ecological and field data.

## Acknowledgements

We thank Isabella Tyler and Hayden Elliott for their support with mosquito rearing at the Department of Entomology, Texas A&M University; Susan Bennett at the Center for Vector-Borne Infectious Diseases at Colorado State University, for kindly providing *Culex quinquefasciatus* and *Culex tarsalis* eggs; and the World Reference Center for Viruses and Arboviruses at the University of Texas Medical Branch for kindly providing Madariaga virus strain Panama. We also thank the team at the Global Health Research Complex (GHRC) and the Biosafety Office at Texas A&M University for their guidance and support during the optimization of the Biosafety Level 3 protocols and execution of experiments at the GHRC. This work was funded by Texas A&M AgriLife Research (including a FY24-25 Insect Vectored Diseases Seed grant granted to T. Magalhaes), and the United States Department of Agriculture National Institute of Food and Agriculture (Hatch # 6070).

## Supplementary Tables

**Suppl. Table 1:** Infection probabilities of body, legs and saliva collected from different mosquito species exposed to Madariaga virus (strain Panama). Least-squares means of infection probabilities and 95% confidence intervals were estimated using logistic regression models.

| Mosquito species | Infection probability [95% CI] |  |  |
| --- | --- | --- | --- |
|  | Body | Legs | Saliva |
| *Aedes aegypti* | 0.622 [0.461-0.759] | 0.328 [0.2-0.489] | 0.201 [0.102-0.357] |
| *Aedes albopictus* | 0.297 [0.175-0.457] | 0.161 [0.077-0.306] | 0.1 [0.038-0.242] |
| *Aedes taeniorhynchus* | 0.412 [0.213-0.645] | 0.345 [0.157-0.598] | 0.104 [0.025-0.349] |
| *Culex coronator* | 0.23 [0.118-0.4] | 0.151 [0.068-0.301] | 0.133 [0.052-0.303] |
| *Culex tarsalis* | 0.334 [0.226-0.462] | 0.302 [0.199-0.431] | 0.116 [0.056-0.226] |
| *Culex quinquefasciatus* * | 0.15 [0.08-0.265] | 0.033 [0.008-0.124] | - |

\**Culex quinquefasciatus* had no positive saliva and was excluded from the saliva model. PFU/mL: plaque forming units per mL. 95% CI: 95% confidence interval. Pairwise comparisons were performed through odds ratio analysis.

**Supp. Table 2:**
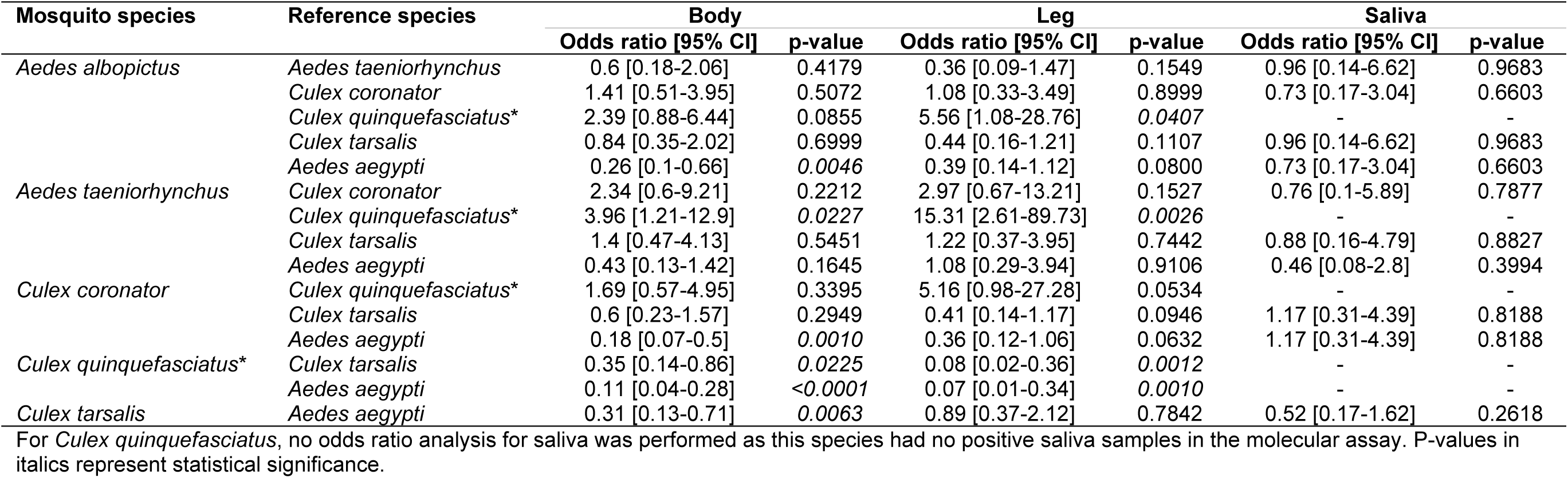
Odds ratios derived from least squares means of body, legs, and saliva infection probabilities in mosquitoes infected with Madariaga virus (strain Panama).

**Supp. Table 3:** Pairwise comparisons of Madariaga virus (strain Panama) infection probability in saliva samples between *Culex quinquefasciatus* and other mosquito species. Samples were collected at 14 days-post exposure.

| Species 1 | Species 2 | Fisher's exact test |  |  |
| --- | --- | --- | --- | --- |
|  |  | Raw p-value | Bonferroni-adjusted p-value | FDR-adjusted p-value |
| *Culex quinquefasciatus* | *Aedes aegypti* | *0.0004* | *0.0020* | *0.0020* |
|  | *Aedes albopictus* | *0.0233* | 0.1165 | *0.0291* |
|  | *Aedes taeniorhynchus* | *0.0299* | 0.1495 | *0.0299* |
|  | *Culex coronator* | *0.0074* | *0.0370* | *0.0185* |
|  | *Culex tarsalis* | *0.0130* | 0.0650 | *0.0217* |

P-values in italics represent statistical significance.

**Supp. Table 4:**
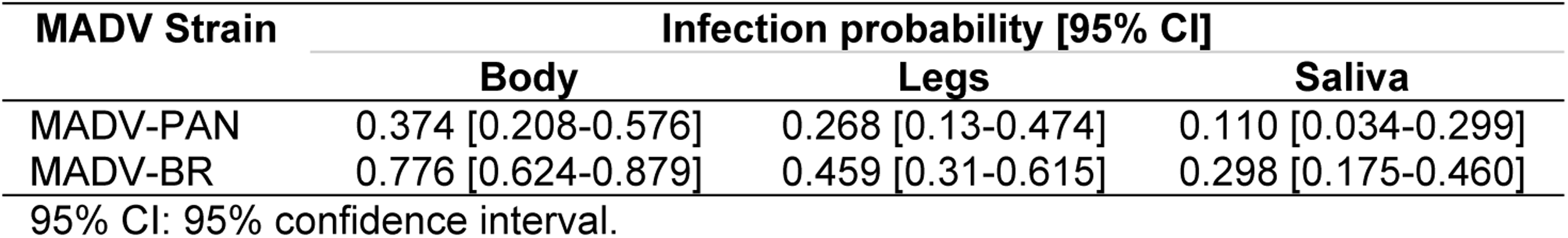
Infection probabilities of body, legs and saliva collected from *Aedes taeniorhynchus* exposed to Madariaga strain Panama (MADV-PAN) or Madariaga virus strain Brazil (MADV-BR). Least-squares means of infection probabilities and 95% confidence intervals were estimated using logistic regression models.

**Supp. Table 5:**
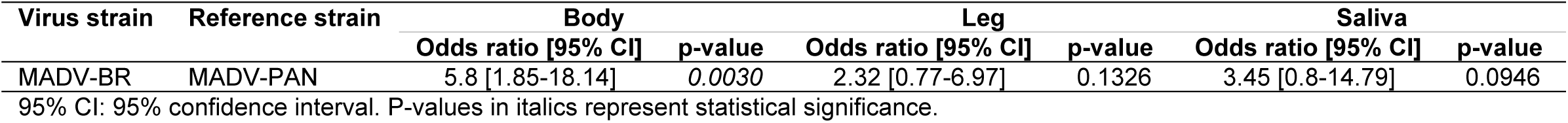
Odds ratios derived from least squares means of the infection probabilities of Madariaga virus strain Panama (MADV-PAN) and Madariaga virus strain Brazil (MADV-BR), in body, legs and saliva collected from *Aedes taeniorhynchus* at 14 days-post exposure.

**Supp. Table 6:** Infection probabilities of body, legs and saliva collected from *Aedes aegypti and Aedes* albopictus infected with Madariaga (strain Panama), at 3, 7, 14, and 21 days-post exposure (dpe). Least-squares means of infection probabilities and 95% confidence intervals were estimated using logistic regression models.

| Mosquito species | Dpe | Infection probability [95% CI] |  |  |
| --- | --- | --- | --- | --- |
|  |  | Body | Legs | Saliva |
| *Aedes aegypti* | 3 | 0.325 [0.198-0.483] | 0.05 [0.012-0.179] | 0.0 [0.0-1.0] |
|  | 7 | 0.325 [0.198-0.483] | 0.124 [0.052-0.267] | 0.049 [0.012-0.178] |
|  | 14 | 0.625 [0.467-0.761] | 0.349 [0.218-0.508] | 0.198 [0.102-0.351] |
|  | 21 | 0.826 [0.676-0.915] | 0.5 [0.349-0.651] | 0.324 [0.197-0.483] |
| *Aedes albopictus* | 3 | 0.274 [0.158-0.432] | 0.274 [0.158-0.432] | 0.049 [0.012-0.178] |
|  | 7 | 0.65 [0.492-0.781] | 0.274 [0.158-0.432] | 0.149 [0.068-0.295] |
|  | 14 | 0.299 [0.178-0.458] | 0.174 [0.085-0.324] | 0.099 [0.037-0.237] |
|  | 21 | 0.1 [0.038-0.238] | 0.05 [0.012-0.179] | 0.0 [0.0-1.0] |

95% CI: 95% confidence interval.

**Supp. Table 7:** Odds ratios derived from least squares means of body, legs, and saliva infection probabilities in *Aedes aegypti* and *Aedes albopictus* infected with Madariaga virus (strain Panama), at 3, 7, 14, and 21 days-post exposure (dpe).

| Mosquito species | Reference species | Dpe | Body |  | Legs |  | Saliva |  |
| --- | --- | --- | --- | --- | --- | --- | --- | --- |
|  |  |  | Odds ratio [95% CI] | p-value | Odds ratio [95% CI] | p-value | Odds ratio [95% CI] | p-value |
| *Aedes albopictus* | *Aedes aegypti* | 3 | 0.79 [0.3-2.06] | 0.6258 | 7.23 [1.48-35.44] | 0.0149 | - | - |
|  |  | 7 | 3.87 [1.52-9.84] | 0.0046 | 2.66 [0.82-8.59] | 0.1013 | 3.36 [0.63-17.95] | 0.1550 |
|  |  | 14 | 0.26 [0.1-0.65] | 0.0045 | 0.39 [0.14-1.12] | 0.0805 | 0.44 [0.12-1.62] | 0.2183 |
|  |  | 21 | 0.02 [0.01-0.09] | <0.0001 | 0.05 [0.01-0.25] | 0.0002 | - | - |

95% CI: 95% confidence interval. *Aedes aegypti* at 3 dpe and *Ae. albopictus* at 21 dpe had no positive saliva samples, therefore the odds ratio was not reliably estimated.

**Supp. Table 8:** Mean log_10_-transformed plaque forming units per mL (PFU/mL) of Madariaga virus (strain Panama), in body, leg and saliva samples collected from six mosquito species at 14 days-post infection.

| Mosquito species | Mean Log<sub>10</sub> MADV-PAN PFU/mL [95% CI] |  |  |
| --- | --- | --- | --- |
|  | Body | Legs | Saliva |
| *Aedes aegypti* | 5.51 [5.39-5.64] | 5.46 [5.33-5.58] | 3.93 [3.77-4.10] |
| *Aedes albopictus* | 5.90 [5.69-6.10] | 5.13 [4.93-5.33] | 3.74 [3.54-3.94] |
| *Aedes taeniorhynchus* | 5.76 [5.55-5.97] | 5.49 [5.30-5.68] | 4.29 [3.76-4.83] |
| *Culex coronator* | 5.33 [5.18-5.48] | 5.23 [5.09-5.38] | 4.05 [3.82-4.27] |
| *Culex tarsalis* | 5.18 [5.06-5.31] | 5.39 [5.27-5.51] | 4.15 [4.01-4.29] |
| *Culex quinquefasciatus* * | 5.22 [5.08-5.36] | 5.34 [5.08-5.59] | - |

95% CI: 95% confidence interval. \**Culex quinquefasciatus* did not have any positive saliva samples in the molecular assay.

**Supp. Table 9:** Mean log_10_-transformed plaque forming units per mL (PFU/mL) of Madariaga virus strain Panama (MADV-PAN) or Madariaga virus strain Brazil (MADV-BR), in body, leg and saliva samples collected from *Aedes taeniorhynchus* at 14 days-post infection.

| Mosquito species | Virus lineage | Mean Log<sub>10</sub> MADV PFU/mL [95% CI] |  |  |
| --- | --- | --- | --- | --- |
|  |  | Body | Legs | Saliva |
| *Aedes taeniorhynchus* | MADV-PAN | 5.75 [5.62-5.88] | 5.49 [5.35-5.63] | 4.08 [3.75-4.42] |
|  | MADV-BR | 5.34 [5.26-5.42] | 5.16 [5.07-5.26] | 3.78 [3.62-3.94] |

95% CI: 95% confidence interval.

**Supp. Table 10:**
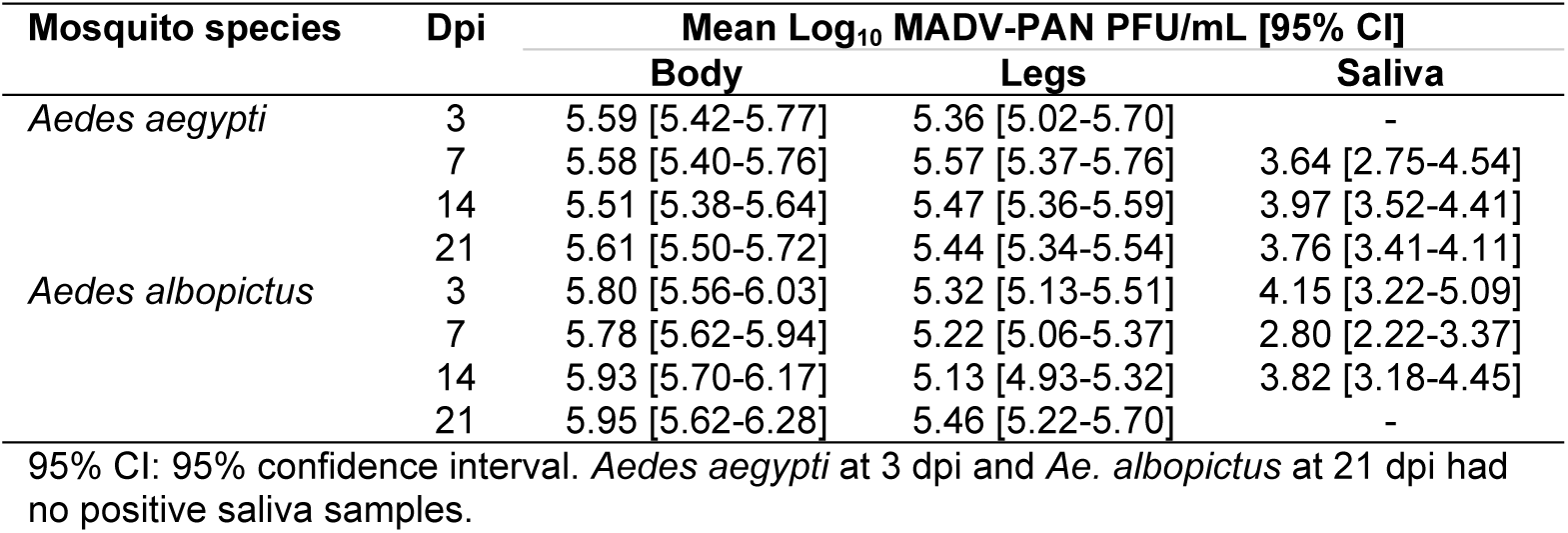
Mean log_10_-transformed plaque forming units per mL (PFU/mL) of Madariaga virus strain Panama (MADV-PAN), in body, leg and saliva samples collected from *Aedes aegypti* and *Aedes albopictus* at 3, 7, 14, and 21 days-post infection.

| Mosquito species | Dpi | Mean Log<sub>10</sub> MADV-PAN PFU/mL [95% CI] |  |  |
| --- | --- | --- | --- | --- |
|  |  | Body | Legs | Saliva |
| *Aedes aegypti* | 3 | 5.59 [5.42-5.77] | 5.36 [5.02-5.70] | - |
|  | 7 | 5.58 [5.40-5.76] | 5.57 [5.37-5.76] | 3.64 [2.75-4.54] |
|  | 14 | 5.51 [5.38-5.64] | 5.47 [5.36-5.59] | 3.97 [3.52-4.41] |
|  | 21 | 5.61 [5.50-5.72] | 5.44 [5.34-5.54] | 3.76 [3.41-4.11] |
| *Aedes albopictus* | 3 | 5.80 [5.56-6.03] | 5.32 [5.13-5.51] | 4.15 [3.22-5.09] |
|  | 7 | 5.78 [5.62-5.94] | 5.22 [5.06-5.37] | 2.80 [2.22-3.37] |
|  | 14 | 5.93 [5.70-6.17] | 5.13 [4.93-5.32] | 3.82 [3.18-4.45] |
|  | 21 | 5.95 [5.62-6.28] | 5.46 [5.22-5.70] | - |

95% CI: 95% confidence interval. *Aedes aegypti* at 3 dpi and *Ae. albopictus* at 21 dpi had no positive saliva samples.

## References

1. Magalhaes T, Hamer GL, de Carvalho-Leandro D, Ribeiro VML, Turell MJ. Uncertainties surrounding Madariaga virus, a member of the eastern equine encephalitis virus complex. Vector Borne Zoonotic Dis. 2024 Oct;24(10):633–40.

2. Alice FJ. Infecção humana pelo vírus “leste” da encefalite equina. Bol Inst Biol Bahia. 1956;3:3–9.

3. Corniou B, Ardoin P, Bartholomew C, Ince W, Massiah V. First isolation of a South American strain of eastern equine virus from a case of encephalitis in Trinidad. Trop Geogr Med. 1972 Jun;24(2):162–7.

4. Carrera JP, Forrester N, Wang E, Vittor AY, Haddow AD, Lopez-Verges S, et al. Eastern equine encephalitis in Latin America. N Engl J Med. 2013 Aug 22;369(8):732–44.

5. Carrera JP, Cucunuba ZM, Neira K, Lambert B, Pitti Y, Liscano J, et al. Endemic and epidemic human alphavirus infections in eastern Panama: an analysis of population-based cross-sectional surveys. Am J Trop Med Hyg. 2020 Dec;103(6):2429–37.

6. Rivera LF, Lezcano-Coba C, Galue J, Rodriguez X, Juarez Y, de Souza WM, et al. Characteristics of Madariaga and Venezuelan equine encephalitis virus infections, Panama. Emerg Infect Dis. 2024;30(14):94–104.

7. Shope RE, de Andrade AH, Bensabath G, Causey OR, Humphrey PS. The epidemiology of EEE WEE, SLE and Turlock viruses, with special reference to birds, in a tropical rain forest near Belem, Brazil. Am J Epidemiol. 1966 Nov;84(3):467–77.

8. Srihongse S, Galindo P. The isolation of eastern equine encephalitis from *Culex* (*Melanoconion*) *taeniopus* Dyar and Knab in Panama. Mosquito News. 1967;27(1):3.

9. Dietz WH, Jr., Galindo P, Johnson KM. Eastern equine encephalomyelitis in Panama: the epidemiology of the 1973 epizootic. Am J Trop Med Hyg. 1980 Jan;29(1):133–40.

10. Walder R, Suarez OM, Calisher CH. Arbovirus studies in the Guajira region of Venezuela: activities of eastern equine encephalitis and Venezuelan equine encephalitis viruses during an interepizootic period. Am J Trop Med Hyg. 1984 Jul;33(4):699–707.

11. Vasconcelos PFC, Travassos da Rosa JFS, Travassos da Rosa APA, Degallier N, Pinheiro FP, Sá Filho GC. Epidemiologia das encefalites por arbovírus na amazônia brasileira. Rev Inst Med Trop S Paulo. 1991;33(6):465–76.

12. Turell MJ, O’Guinn ML, Jones JW, Sardelis MR, Dohm DJ, Watts DM, et al. Isolation of viruses from mosquitoes (Diptera: Culicidae) collected in the Amazon Basin region of Peru. J Med Entomol. 2005 Sep;42(5):891–8.

13. Causey OR, Causey CE, Maroja OM, Macedo DG. The isolation of arthropod-borne viruses, including members of two hitherto undescribed serological groups, in the Amazon region of Brazil. Am J Trop Med Hyg. 1961 Mar;10:227–49.

14. Turell MJ, O’Guinn ML, Dohm D, Zyzak M, Watts D, Fernandez R, et al. Susceptibility of Peruvian mosquitoes to eastern equine encephalitis virus. J Med Entomol. 2008 Jul;45(4):720–5.

15. Gil L, Magalhaes T, Santos B, Oliveira LV, Oliveira-Filho EF, Cunha JLR, et al. Active circulation of Madariaga virus, a member of the eastern equine encephalitis virus complex, in Northeast Brazil. Pathogens. 2021 Aug 3;10(8).

16. Higgs S, Beaty B. Natural cycles of vector-borne pathogens. In: Marquardt WC, Black IV WC, Freier JE, Hagedorn HH, Hemingway J, Higgs S, et al., editors. Biology of disease vectors. 2nd ed. Burlington: Elsevier Academic Press; 2004. p. 785.

17. Turell MJ, Gargan TP, 2nd, Bailey CL. Replication and dissemination of Rift Valley fever virus in *Culex pipiens*. Am J Trop Med Hyg. 1984 Jan;33(1):176–81.

18. Kramer LD, Scherer WF. Vector competence of mosquitoes as a marker to distinguish Central American and Mexican epizootic from enzootic strains of Venezuelan encephalitis virus. Am J Trop Med Hyg. 1976 Mar;25(2):336–46.

19. Rochlin I, Faraji A, Healy K, Andreadis TG. West Nile virus mosquito vectors in North America. J Med Entomol. 2019 Oct 28;56(6):1475–90.

20. Bond JG, Casas-Martinez M, Quiroz-Martinez H, Novelo-Gutierrez R, Marina CF, Ulloa A, et al. Diversity of mosquitoes and the aquatic insects associated with their oviposition sites along the Pacific coast of Mexico. Parasit Vectors. 2014 Jan 22;7:41.

21. Demari-Silva B, Suesdek L, Sallum MA, Marrelli MT. Wing geometry of Culex coronator (Diptera: Culicidae) from south and southeast Brazil. Parasit Vectors. 2014 Apr 9;7:174.

22. Pecor JE, Harbach RE, Peyton EL, Roberts DR, Rejmankova E, Manguin S, et al. Mosquito studies in Belize, Central America: records, taxonomic notes, and a checklist of species. J Am Mosq Control Assoc. 2002 Dec;18(4):241–76.

23. Laurito M, Briscoe AG, Almirón WR, Harbach RE. Systematics of the *Culex coronator* complex (Diptera: Culicidae): morphological and molecular assessment. Zool J Linn Soc. 2018;182:23.

24. Sames WJ, Mann JG, Kelly R, Evans CL, Varnado WC, Bosworth AB, et al. Distribution of *Culex coronator* in the USA. J Am Mosq Control Assoc. 2021 Mar 1;37(1):1–9.

25. Consoli RAGB, Lourenço de Oliveira R. Principais mosquitos de importância sanitária no Brasil. 1st ed. Rio de Janeiro: Editora Fiocruz; 1994.

26. Wilke ABB, Vasquez C, Cardenas G, Carvajal A, Medina J, Petrie WD, et al. Invasion, establishment, and spread of invasive mosquitoes from the Culex coronator complex in urban areas of Miami-Dade County, Florida. Sci Rep. 2021 Jul 16;11(1):14620.

27. Alto BW, Connelly CR, O’Meara GF, Hickman D, Karr N. Reproductive biology and susceptibility of Florida *Culex coronator* to infection with West Nile virus. Vector Borne Zoonotic Dis. 2014 Aug;14(8):606–14.

28. Scott TW, Takken W. Feeding strategies of anthropophilic mosquitoes result in increased risk of pathogen transmission. Trends Parasitol. 2012 Mar;28(3):114–21.

29. Cebrian-Camison S, Martinez-de la Puente J, Figuerola J. A literature review of host feeding patterns of invasive *Aedes* mosquitoes in Europe. Insects. 2020 Nov 29;11(12).

30. Olson MF, Ndeffo-Mbah ML, Juarez JG, Garcia-Luna S, Martin E, Borucki MK, et al. High rate of non-human feeding by *Aedes aegypti* reduces Zika virus transmission in South Texas. Viruses. 2020 Apr 17;12(4).

31. Agha SB, Tchouassi DP, Turell MJ, Bastos ADS, Sang R. Entomological assessment of dengue virus transmission risk in three urban areas of Kenya. PLoS Negl Trop Dis. 2019 Aug;13(8):e0007686.

32. Garcia-Rejon JE, Navarro JC, Cigarroa-Toledo N, Baak-Baak CM. An updated review of the invasive *Aedes albopictus* in the Americas; geographical distribution, host feeding patterns, arbovirus infection, and the potential for vertical transmission of dengue virus. Insects. 2021 Oct 26;12(11).

33. Kraemer MU, Sinka ME, Duda KA, Mylne AQ, Shearer FM, Barker CM, et al. The global distribution of the arbovirus vectors *Aedes aegypti* and *Ae. albopictus*. Elife. 2015 Jun 30;4:e08347.

34. Pereira-Dos-Santos T, Roiz D, Lourenco-de-Oliveira R, Paupy C. A systematic review: is *Aedes albopictus* an efficient bridge vector for zoonotic arboviruses? Pathogens. 2020 Apr 7;9(4).

35. Gutierrez-Lopez R, Bialosuknia SM, Ciota AT, Montalvo T, Martinez-de la Puente J, Gangoso L, et al. Vector competence of *Aedes caspius* and *Ae. albopictus* mosquitoes for Zika virus, Spain. Emerg Infect Dis. 2019 Feb;25(2):346–8.

36. Obadia T, Gutierrez-Bugallo G, Duong V, Nunez AI, Fernandes RS, Kamgang B, et al. Zika vector competence data reveals risks of outbreaks: the contribution of the European ZIKAlliance project. Nat Commun. 2022 Aug 2;13(1):4490.

37. Roundy CM, Azar SR, Rossi SL, Huang JH, Leal G, Yun R, et al. Variation in *Aedes aegypti* mosquito competence for Zika virus transmission. Emerg Infect Dis. 2017 Apr;23(4):625–32.

38. Salazar MI, Richardson JH, Sanchez-Vargas I, Olson KE, Beaty BJ. Dengue virus type 2: replication and tropisms in orally infected *Aedes aegypti* mosquitoes. BMC Microbiol. 2007 Jan 30;7:9.

39. Turner EA, Clark SD, Pena-Garcia VH, Christofferson RC. Investigating the effects of microclimate on arboviral kinetics in *Aedes aegypti*. Pathogens. 2024 Dec 14;13(12).

40. Anderson JF, Fish D, Armstrong PM, Misencik MJ, Bransfield A, Ferrandino FJ, et al. Seasonal dynamics of mosquito-borne viruses in the Southwestern Florida Everglades, 2016, 2017. Am J Trop Med Hyg. 2022 Jan 10;106(2):610–22.

41. Eastwood G, Goodman SJ, Cunningham AA, Kramer LD. *Aedes taeniorhynchus* vectorial capacity informs a pre-emptive assessment of West Nile virus establishment in Galapagos. Sci Rep. 2013;3:1519.

42. Bataille A, Cunningham AA, Cruz M, Cedeno V, Goodman SJ. Seasonal effects and fine-scale population dynamics of *Aedes taeniorhynchus*, a major disease vector in the Galapagos Islands. Mol Ecol. 2010 Oct;19(20):4491–504.

43. de Melo Ximenes MFF, de Araujo Galvao JM, Inacio CLS, Macedo ESVP, Pereira R, Pinheiro MPG, et al. Arbovirus expansion: New species of culicids infected by the Chikungunya virus in an urban park of Brazil. Acta Trop. 2020 Sep;209:105538.

44. Dos Reis IC, Gibson G, Ayllon T, de Medeiros Tavares A, de Araujo JMG, da Silva Monteiro E, et al. Entomo-virological surveillance strategy for dengue, Zika and chikungunya arboviruses in field-caught *Aedes* mosquitoes in an endemic urban area of the Northeast of Brazil. Acta Trop. 2019 Sep;197:105061.

45. Ryba J, Fuentes O, Danielova V, Fernandez A. Mosquito studies on the Isla de la Juventud, Cuba, at the beginning of rain period. Folia Parasitol (Praha). 1984;31(2):163–7.

46. Scherer WF, Dickerman RW, Ordonez JV, Seymour C, 3rd, Kramer LD, Jahrling PB, et al. Ecologic studies of Venezuelan encephalitis virus and isolations of Nepuyo and Patois viruses during 1968-1973 at a marsh habitat near the epicenter of the 1969 outbreak in Guatemala. Am J Trop Med Hyg. 1976 Jan;25(1):151–62.

47. Asigau S, Salah S, Parker PG. Assessing the blood meal hosts of *Culex quinquefasciatus* and *Aedes taeniorhynchus* in Isla Santa Cruz, Galapagos. Parasit Vectors. 2019 Dec 16;12(1):584.

48. Ortiz YV, Casas SA, Tran MND, Decker EG, Saborit I, Le HN, et al. Mosquito population dynamics and blood host associations in two types of urban greenspaces in coastal Florida. Insects. 2025 Feb 20;16(3).

49. Turell MJ. Effect of environmental temperature on the vector competence of *Aedes taeniorhynchus* for Rift Valley fever and Venezuelan equine encephalitis viruses. Am J Trop Med Hyg. 1993 Dec;49(6):672–6.

50. Turell MJ. Vector competence of three Venezuelan mosquitoes (Diptera: Culicidae) for an epizootic IC strain of Venezuelan equine encephalitis virus. J Med Entomol. 1999 Jul;36(4):407–9.

51. Turell MJ, Beaman JR, Neely GW. Experimental transmission of eastern equine encephalitis virus by strains of *Aedes albopictus* and *A. taeniorhynchus* (Diptera: Culicidae). J Med Entomol. 1994 Mar;31(2):287–90.

52. Turell MJ, Ludwig GV, Beaman JR. Transmission of Venezuelan equine encephalomyelitis virus by *Aedes sollicitans* and *Aedes taeniorhynchus* (Diptera: Culicidae). J Med Entomol. 1992 Jan;29(1):62–5.

53. Turell MJ, O’Guinn ML, Dohm DJ, Jones JW. Vector competence of North American mosquitoes (Diptera: Culicidae) for West Nile virus. J Med Entomol. 2001 Mar;38(2):130–4.

54. Turell MJ, Rossi CA, Bailey CL. Effect of extrinsic incubation temperature on the ability of *Aedes taeniorhynchus* and *Culex pipiens* to transmit Rift Valley fever virus. Am J Trop Med Hyg. 1985 Nov;34(6):1211–8.

55. Yuill TM, Thompson PH. Cache Valley virus in the Del Mar Va Peninsula. IV. Biological transmission of the virus by *Aedes sollicitans* and *Aedes taeniorhynchus*. Am J Trop Med Hyg. 1970 May;19(3):513–9.

56. Hodapp CJ, Hillis WD, Dahl EV. Isolation of two arboviruses from *Aedes taeniorhynchus* Wiedemann. J Med Entomol. 1966 Apr;3(1):44–5.

57. Ortiz DI, Wozniak A, Tolson MW, Turner PE, Vaughan DR. Isolation of EEE virus from *Ochlerotatus taeniorhynchus* and *Culiseta melanura* in coastal South Carolina. J Am Mosq Control Assoc. 2003 Mar;19(1):33–8.

58. Belle EA, King SD, Griffiths BB, Grant LS. Epidemiological investigation for arboviruses in Jamaica, West Indies. Am J Trop Med Hyg. 1980 Jul;29(4):667–75.

59. Causey OR, Shope RE, Sutmoller P, Laemmert H. Epizootic eastern equine encephalitis in the Bragança region of Pará, Brazil. Rev Serv Esp Saúde Pública. 1962;12(1):39–45.

